## Supplementary Tables and Figures for "Local adaptation and archaic introgression shape global diversity at human structural variant loci"

| SV ID | SV length | LRS | *P*-value | Ancestry component | Affected gene(s) |
|---|---|---|---|---|---|
| 27407_HG02106_ins | 44 | 124.3 | $2.74 \times 10^{-23}$ | 1 | |
| 1737_HG02106_del | 3490 | 58.5 | $7.69 \times 10^{-9}$ | 1 | *LINC01680* |
| 11915_HG01352_ins | 397 | 45.0 | $7.28 \times 10^{-6}$ | 1 | *KDM2A* |
| **22237_HG02059_ins** | **34** | **513.0** | $\mathbf{5.34 \times 10^{-108}}$ | **2** | ***IGHG4*** |
| **22231_HG02059_del** | **135** | **488.1** | $\mathbf{1.42 \times 10^{-102}}$ | **2** | |
| 25871_AK1_del | 37 | 113.7 | $5.77 \times 10^{-21}$ | 2 | |
| 32021_HG00268_ins | 84 | 98.9 | $1.02 \times 10^{-17}$ | 3 | *PCNT* |
| 2181_CHM1_ins | 309 | 84.6 | $1.36 \times 10^{-14}$ | 3 | *TMEM131* |
| 1731_NA12878_ins | 395 | 82.2 | $4.57 \times 10^{-14}$ | 3 | *LRP1B* |
| 22065_HG02106_del | 186 | 481.1 | $4.67 \times 10^{-101}$ | 4 | |
| 25687_HG02106_ins | 2886 | 468.6 | $2.39 \times 10^{-98}$ | 4 | *CLEC16A* |
| 5843_HG02106_del | 32 | 377.3 | $1.82 \times 10^{-78}$ | 4 | |
| 21859_NA19240_ins | 36 | 159.1 | $6.70 \times 10^{-31}$ | 5 | *AC135050.3* |
| 25014_HG02106_del | 325 | 133.4 | $2.80 \times 10^{-25}$ | 5 | *CSNK1G1, AC087632.2* |
| 21191_NA19240_del | 110 | 122.1 | $8.12 \times 10^{-23}$ | 5 | *AC087632.2, PCLAF* |
| **22237_HG02059_ins** | **34** | **212.2** | $\mathbf{1.73 \times 10^{-42}}$ | **6** | ***IGHG4*** |
| 658_HX1_ins | 445 | 137.6 | $3.42 \times 10^{-26}$ | 6 | |
| **22231_HG02059_del** | **135** | **135.9** | $\mathbf{7.80 \times 10^{-26}}$ | **6** | |
| 10085_HG00268_del | 52 | 116.6 | $1.31 \times 10^{-21}$ | 7 | |
| 18105_NA19240_ins | 3096 | 107.7 | $1.21 \times 10^{-19}$ | 7 | |
| 23087_CHM13_del | 34 | 102.3 | $1.78 \times 10^{-18}$ | 7 | *PSMC3IP* |
| 18075_HG00268_del | 167 | 85.6 | $8.43 \times 10^{-15}$ | 8 | *PLXDC2* |
| 9365_CHM1_ins | 4417 | 83.9 | $1.99 \times 10^{-14}$ | 8 | *CLVS1* |
| 14974_NA19434_ins | 337 | 63.6 | $5.64 \times 10^{-10}$ | 8 | *OR10G3* |

**Table S1.** Highly differentiated SV loci across ancestry components. The top three SVs per ancestry component are reported. The *IGH* insertion and deletion are highlighted in bold text.

| SV ID | Max $r^2$ with Sprime SNP | LRS | Ancestry component | Affected gene(s) |
|---|---|---|---|---|
| 16140_HX1_ins | 1.00 | 86.9 | 6 | |
| 16624_HG00268_del | 1.00 | 53.8 | 3 | |
| 1929_HG02106_del | 1.00 | 173.4 | 4 | *KDM2A* |
| 20771_CHM13_del | 1.00 | 223.1 | 4 | *TCF12* |
| **22231_HG02059_del** | **1.00** | **135.9** | **6** | |
| 24577_HG00514_del | 1.00 | 109.9 | 2 | *TCF25* |
| 29991_HG02106_del | 1.00 | 200.5 | 4 | |
| 3098_HG00268_ins | 1.00 | 50.0 | 3 | |
| 41325_HG04217_del | 1.00 | 100.3 | 5 | |
| 8362_HG02106_ins | 1.00 | 189.6 | 4 | |
| 19726_CHM13_ins | 0.98 | 177.8 | 4 | |
| 10377_HG02106_ins | 0.97 | 101.1 | 6 | *FBXO38* |
| 16625_HG00268_ins | 0.97 | 53.4 | 3 | |
| 20769_CHM13_ins | 0.96 | 225.8 | 4 | *TCF12* |
| 18686_HG01352_ins | 0.96 | 77.1 | 6 | |
| 10847_AK1_del | 0.96 | 81.2 | 3 | *AC007000.4* |
| 12550_HG00733_ins | 0.94 | 36.8 | 1 | *LINC00301* |
| 10373_HG02106_ins | 0.92 | 171.6 | 4 | |
| **22231_HG02059_del** | **0.88** | **488.1** | **2** | |
| 7453_HG02059_ins | 0.83 | 74.1 | 2 | |
| **22237_HG02059_ins** | **0.78** | **513.0** | **2** | ***IGHG4*** |
| 22237_HG02059_ins | 0.78 | 513.0 | 6 | *IGHG4* |
| 12546_HG00733_ins | 0.78 | 32.7 | 1 | *MS4A19P* |
| 32432_HG00268)del | 0.67 | 153.9 | 4 | |
| **22237_HG02059_ins** | **0.64** | **212.2** | **6** | ***IGHG4*** |
| 24574_HG00514_ins | 0.62 | 95.0 | 2 | *SPIRE2* |
| 6305_CHM13_del | 0.58 | 91.5 | 6 | *TLR1* |
| 13529_AK1_ins | 0.55 | 71.6 | 6 | |
| 5444_HG00514_ins | 0.51 | 88.4 | 2 | *NEPRO* |

**Table S2.** Highly differentiated SVs in LD ($r^2 > 0.5$) with putative archaic introgressed haplotypes called by Sprime. The *IGH* insertion and deletion are highlighted in bold text.

**Supplementary Figures**

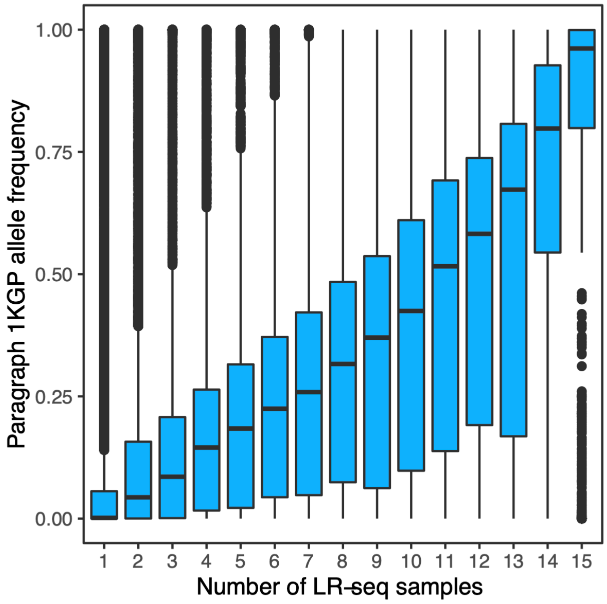


**Figure S1.** Allele frequencies of SVs genotyped with Paragraph in short-read sequenced samples from the 1000 Genomes project compared to the number of PacBio long-read sequenced samples in which the SVs were originally discovered (Audano et al., 2019).

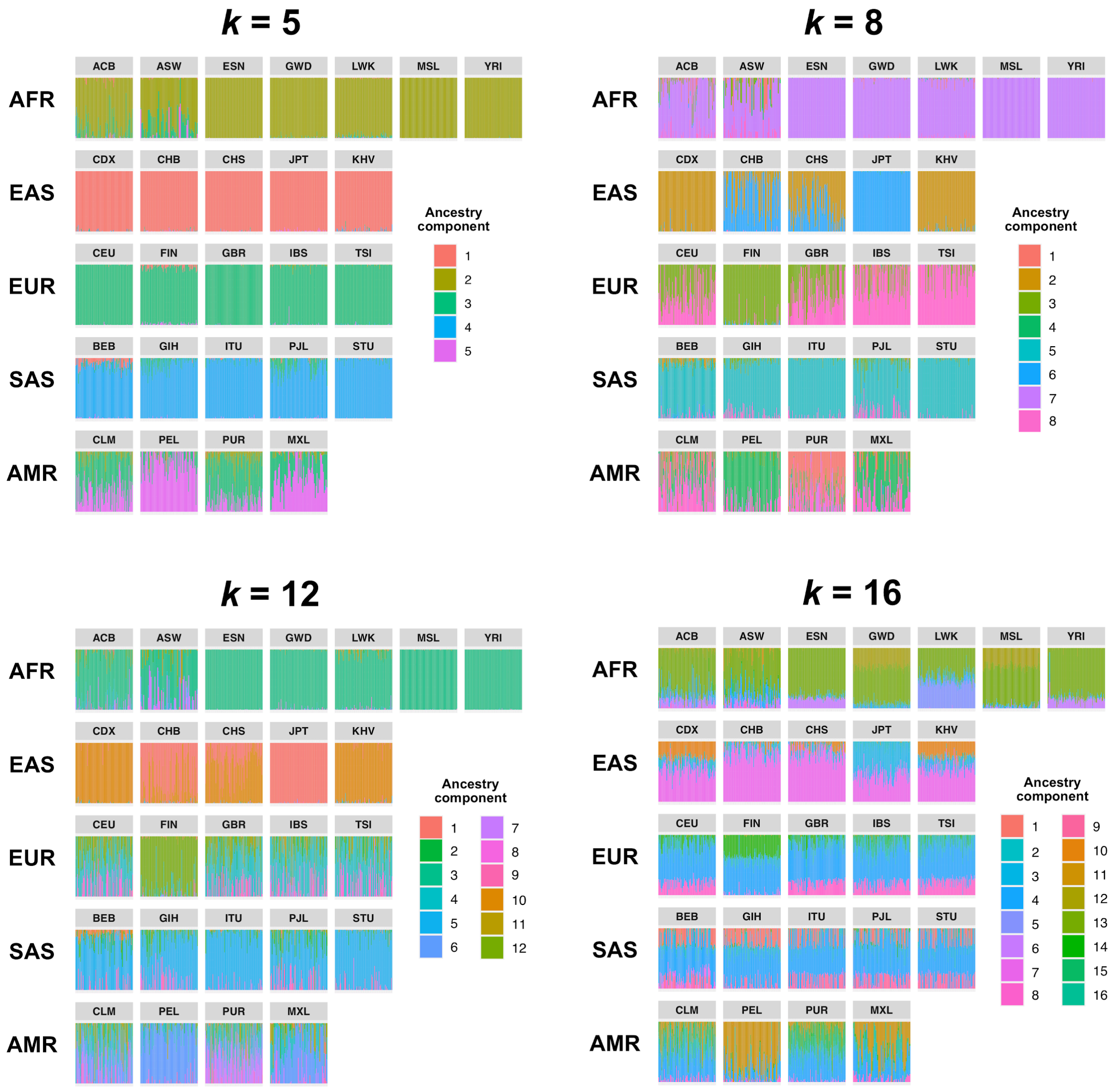


**Figure S2.** Admixture proportions inferred by Ohana for samples in the 1000 Genomes dataset, using *k* = 5, 8 (used in our study), 10, and 16 ancestry components. Vertical bars represent individual samples and are grouped by population.

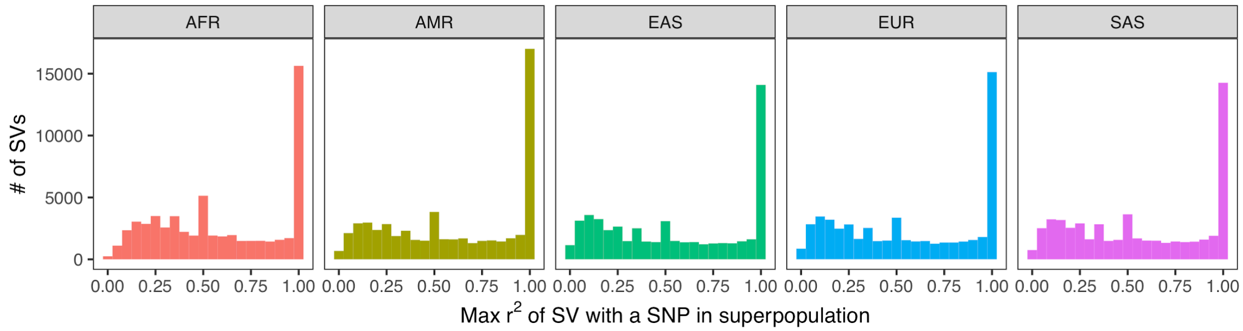


**Figure S3.** Histogram of LD between SVs and neighboring SNPs and short indels. LD was calculated within each of the 26 populations of the 1000 Genomes Project. We then determined the maximum LD of every SV with any of the nearest 100 variants, stratifying by superpopulation.

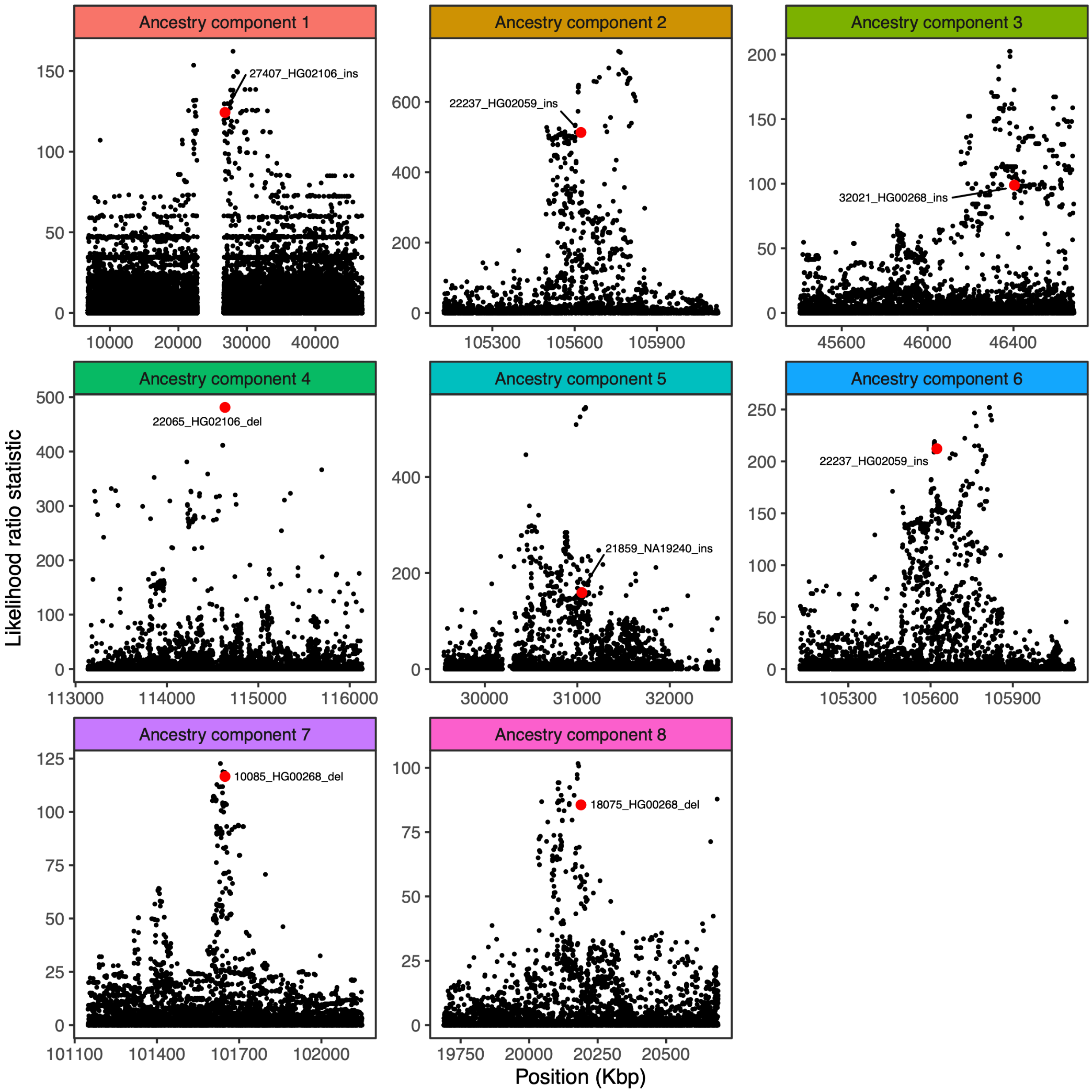


**Figure S4.** Visualization of the likelihood ratio statistic for SNPs and short indels in the genomic vicinity of the top SV hit for each ancestry component. SNPs and short indels are colored in black, while SVs are indicated with red.

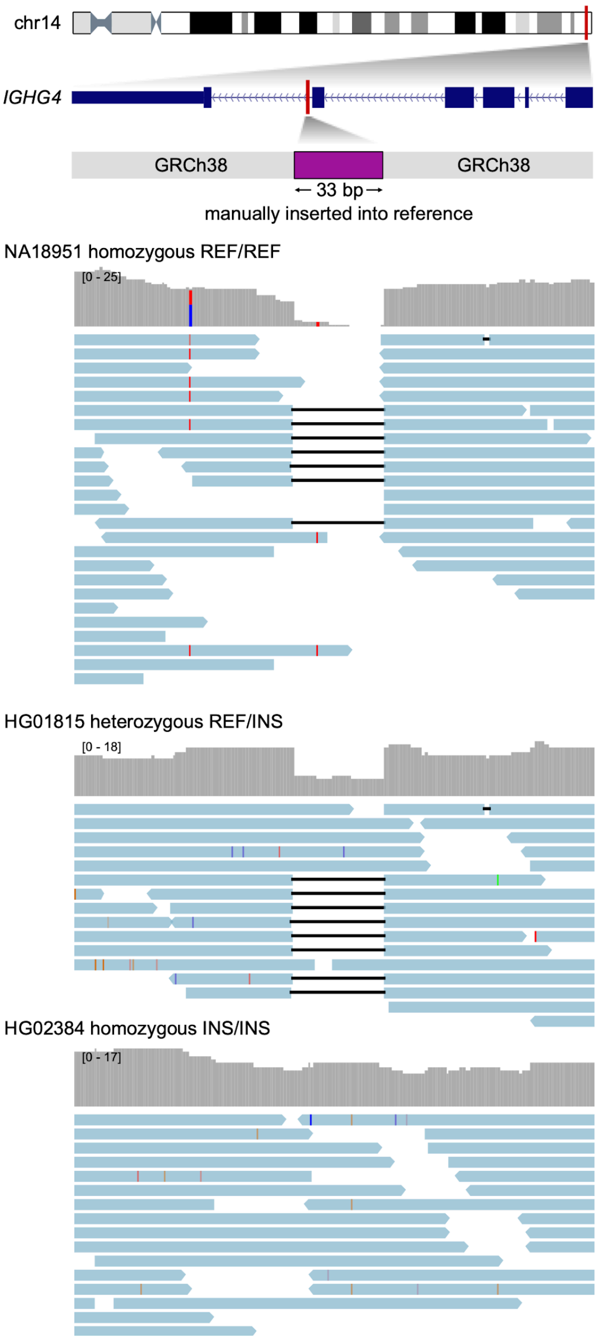


**Figure S5.** Visualization of read alignments to a modified version of the reference genome at the *IGHG4* locus including the intronic 33 bp insertion sequence. Three samples are depicted as representative of each genotype class (homozygous reference, heterozygous, homozygous alternative). Depth of coverage is plotted in upper tracks, while corresponding read alignments are plotted below. Soft-clipped portions of reads were removed to assist visualization.

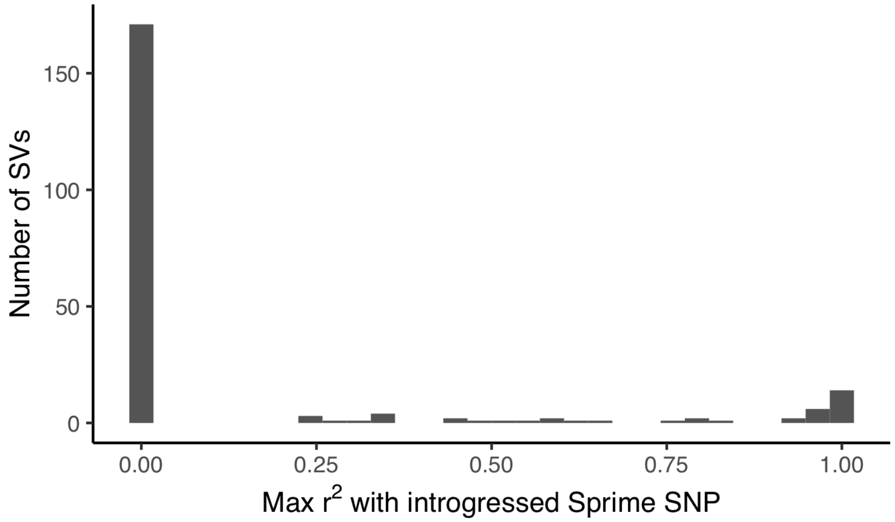


**Figure S6. Distribution of LD between adaptive SVs and introgressed SNPs called by Sprime**. For the 215 most frequency differentiated SVs in our dataset, we calculated LD between the SV and SNPs defining introgressed haplotypes called by Sprime. LD calculations for each SV were restricted to one 1000 Genomes population, chosen based on the ancestry component where the SV was found to exhibit branch-specific differentiation. We identified 26 candidate adaptively introgressed SVs, which had $r^2 > 0.5$ with an introgressed SNP and were at low frequency ($AF < 0.01$) within African populations (excluding admixed ASW and ACB populations).

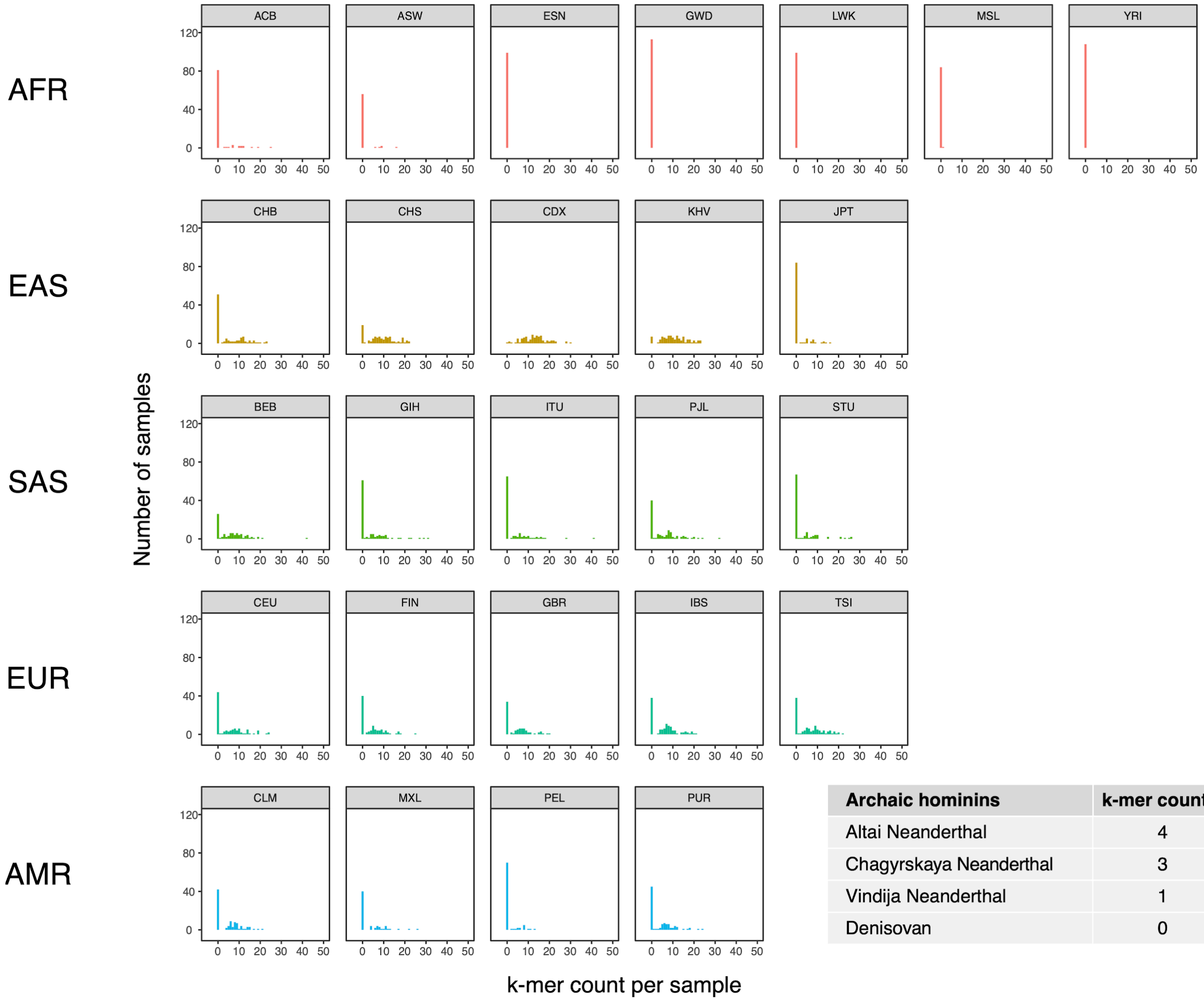


| Archaic hominins | k-mer count |
|---|---|
| Altai Neanderthal | 4 |
| Chagyrskaya Neanderthal | 3 |
| Vindija Neanderthal | 1 |
| Denisovan | 0 |

**Figure S7. Population distribution of *IGHG4* insertion using a diagnostic 48-mer.** Counts of a 48-mer that is diagnostic of the 33 bp *IGHG4* insertion per individual across 1000 Genomes Project dataset, stratified by population. Inset table depicts the counts of the 48-mer in three high-coverage Neanderthal genomes and a high-coverage Denisovan genome.

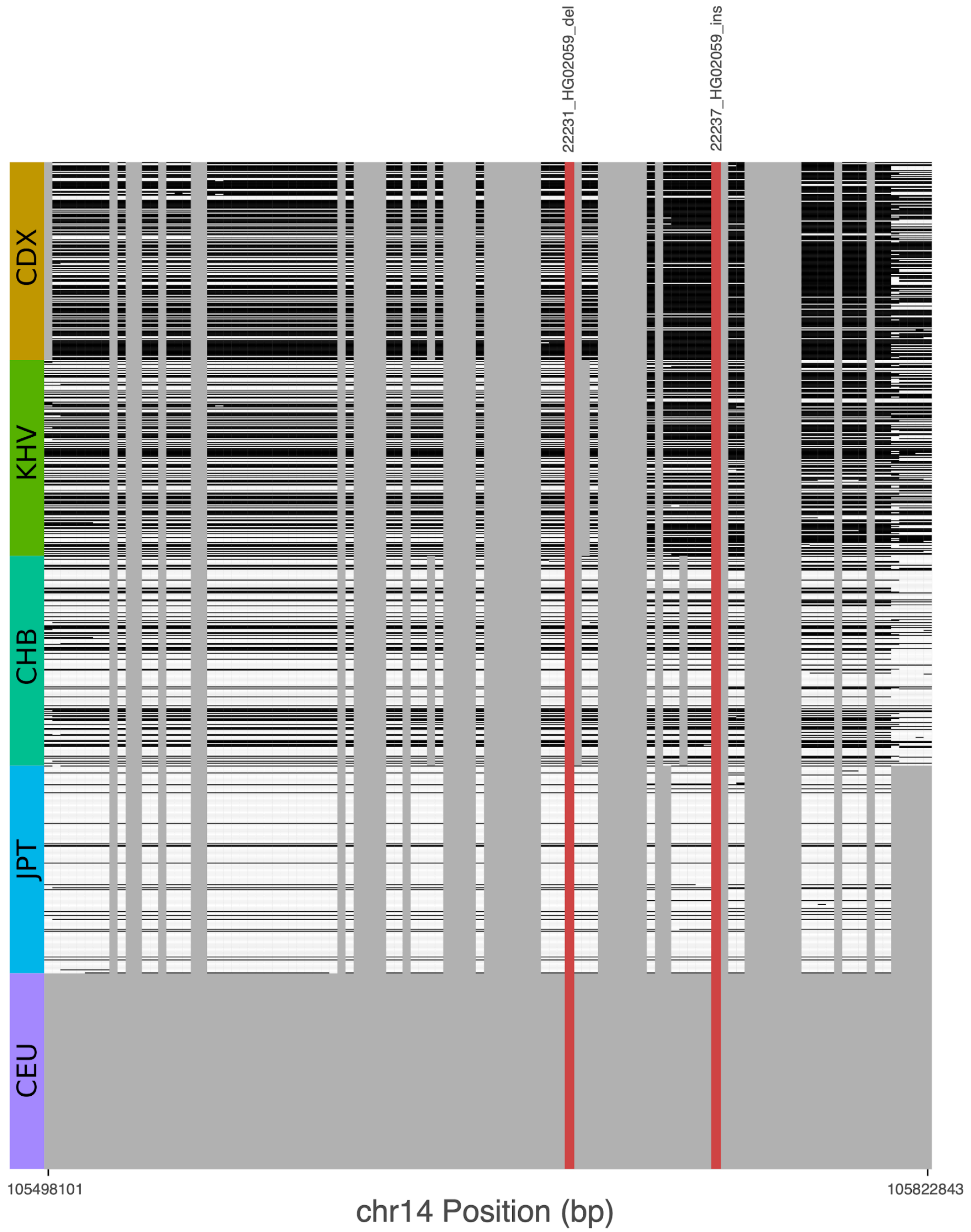


**Figure S8. Signatures of archaic introgression based on calls from Sprime.** For all variants with LRS > 450 at the *IGH* locus, we determined whether these variants fell on Sprime-inferred introgressed haplotypes in individuals of each of five 1000 Genomes project populations. Grey bars represent variants that were not identified by Sprime as introgressed, including short indels that were not included in the analysis by (Browning et al., 2018). Cells are shaded black if the haplotype possesses the putative archaic allele at that variant, and white otherwise. The red bars represent two SVs with the highest LRS in the study.

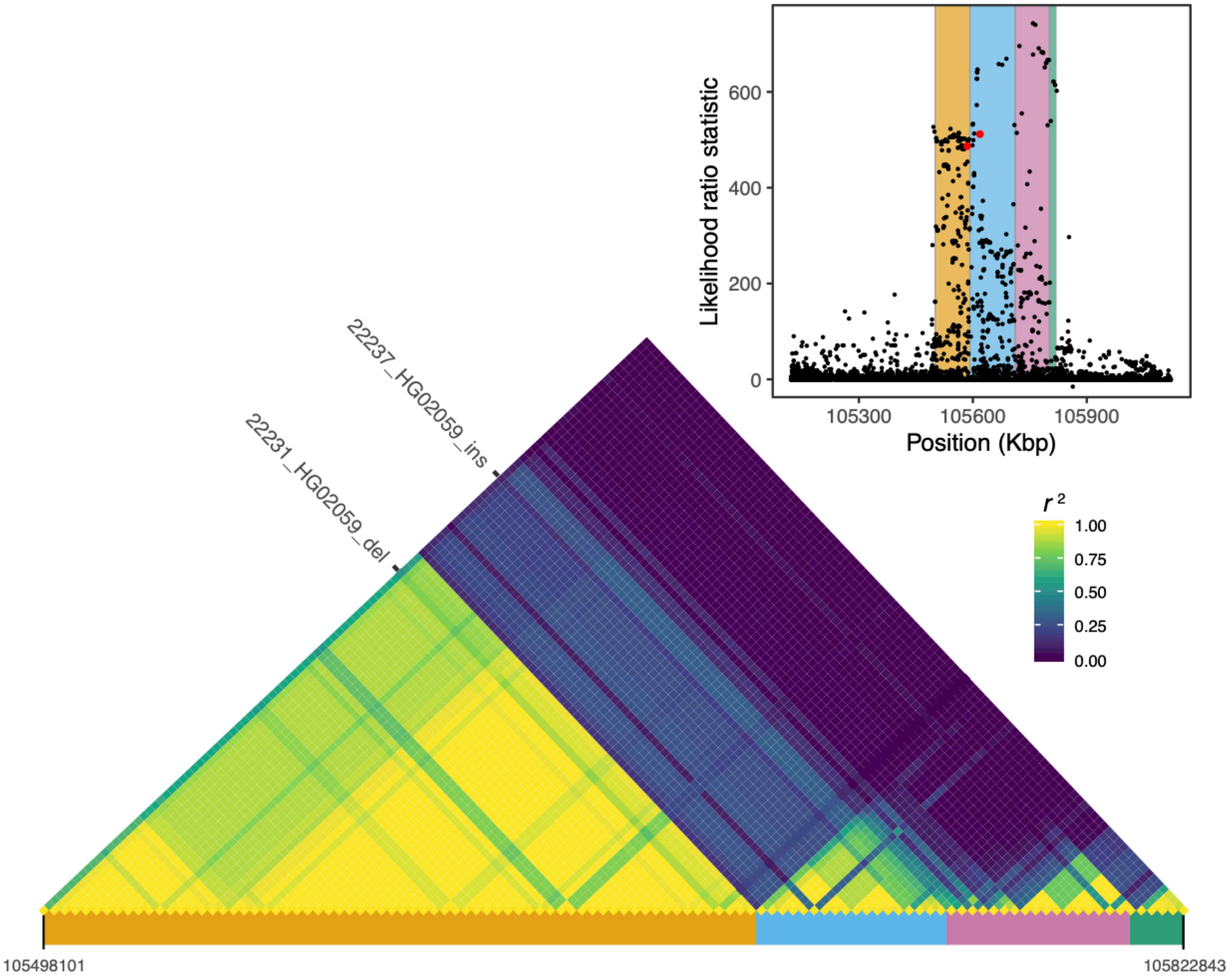


**Figure S9. Local LD at the *IGH* locus.** We calculated pairwise LD between all variants with LRS > 450 at the *IGH* locus, including the two SVs with the highest LRS in our study (22231_HG02059_del, 22237_HG02059_ins; red dots in the inset plot). Visualization of this LD matrix revealed at least four distinct blocks of LD, represented as colored rectangles below the LD heatmap and in the inset plot.

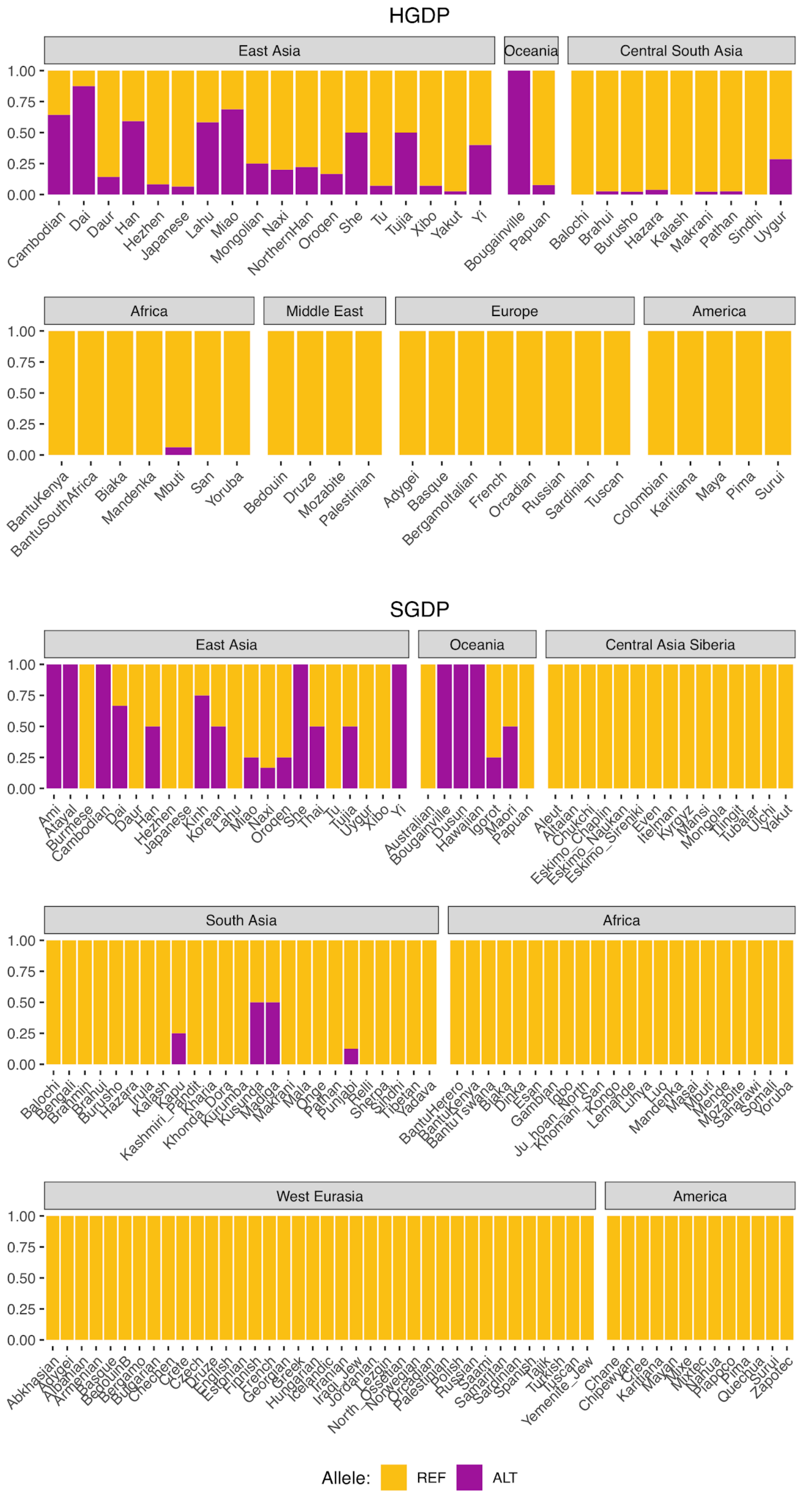


**Figure S10. Global allele frequencies of the introgressed *IGH* haplotype.** Allele frequencies of rs150526114, a SNP that tags the *IGH* haplotype and *IGHG4* insertion, based on data from the Human Genome Diversity Project (HGDP) and the Simons Genome Diversity Project (SGDP).

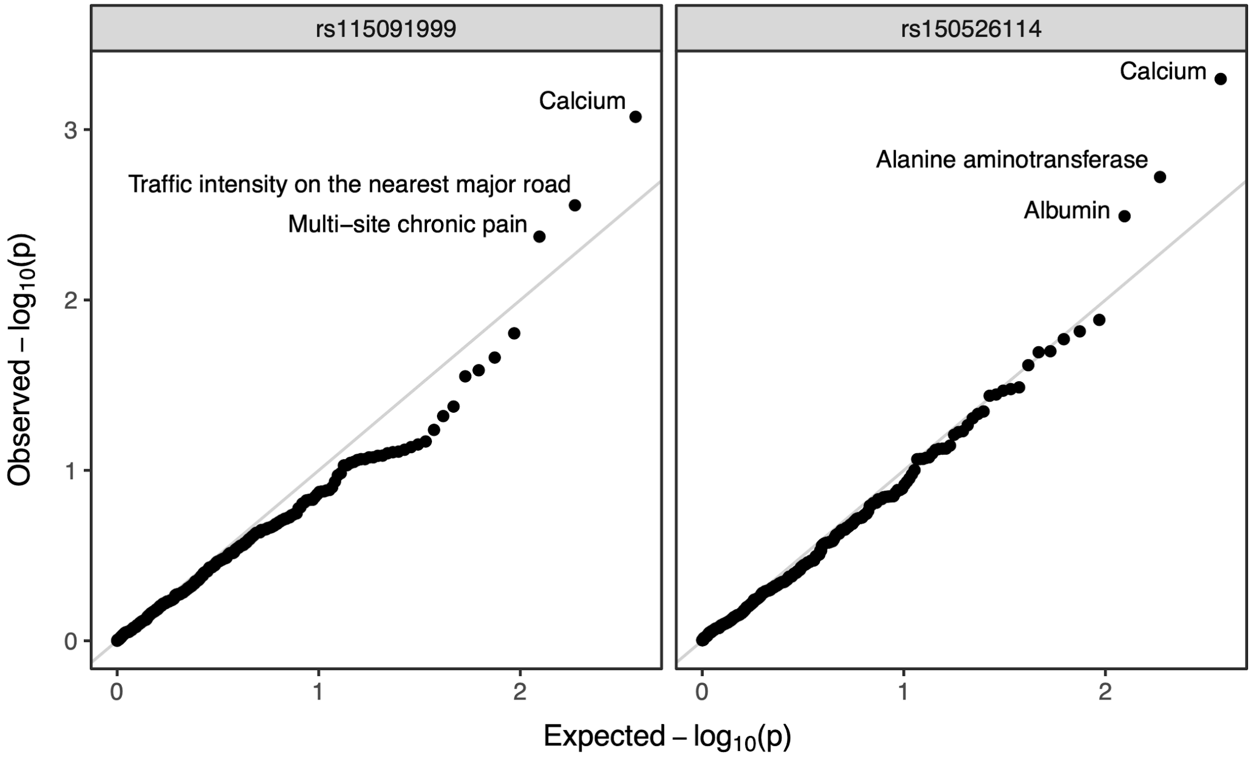


**Figure S11.** Annotated Q-Q plot from a phenome-wide association analysis of putative Neanderthal-introgressed variants at the IGH locus using summary statistics from East Asian individuals from the pan-ancestry GWAS of the UK Biobank dataset. SNP rs115091999 (left panel) tags 22231_HG02059_del, while rs150526114 (right panel) tags 22237_HG02059_ins within the CDX population. No phenotype association is significant after multiple testing correction (Bonferroni-adjusted p-value > 0.05).
